# Biliatresone treatment of pregnant mice causes changes in bile metabolism and liver inflammation in their offspring

**DOI:** 10.1101/2023.03.02.530913

**Authors:** Kapish Gupta, Jimmy P. Xu, Tamir Diamond, Iris De Jong, Andrew Glass, Jessica Llewellyn, Neil D. Theise, Jeffrey D. Winkler, Edward M. Behrens, Clementina Mesaros, Rebecca G. Wells

## Abstract

**Background & Aims:** Biliary atresia is a neonatal disease characterized by bile duct and liver damage, fibrosis, inflammation and abnormal bile metabolism. It appears to result from a prenatal exposure that spares the mother and affects the fetus. Our aim was to define the phenotype in neonatal mice after maternal exposure to low-dose biliatresone, a plant toxin implicated in biliary atresia in livestock.

**Methods:** Pregnant mice were treated orally with low-doses of biliatresone. Histological changes, bile acid profiles and immune profiles were analyzed in postnatal day 5 and 21 pups born to treated mothers.

**Results:** The pups of mothers treated with this dose of biliatresone had no evidence of significant liver or ductular injury or fibrosis at postnatal day 5 or 21 and they grew normally. However, serum levels of glycocholic acid were elevated at postnatal day 5, suggesting altered bile metabolism, and bile metabolism became increasingly abnormal through postnatal day 21, with enhanced glycine conjugation of bile acids. There was also immune cell activation observed in the liver at postnatal day 21.

**Conclusion:** Prenatal exposure to low doses of an environmental toxin can cause liver inflammation and aberrant bile metabolism even in the absence of histological changes.

**Lay summary:** Prenatal exposure to low doses of an environmental toxin can cause changes in bile metabolism in neonatal mice.

## Introduction

Biliary atresia (BA) is a fibrosing neonatal cholangiopathy occurring worldwide with an incidence ranging from 1 in 3,500 to 1 in 18,000 live births [1]. Clinically, BA is associated with extrahepatic bile duct (EHBD) obstruction and fibrosis, bile duct inflammation, and ultimately liver fibrosis and cirrhosis. Additionally, systemic levels of bile acids are elevated [2–4]. However, it is not known whether bile metabolism is altered independently of EHBD obstruction and the resulting abnormal enterohepatic circulation and whether it precedes or results from EHBD obstruction.

BA is thought to result from a prenatal environmental insult, potentially a viral infection or toxic exposure [1]. We have shown that biliatresone, a toxin implicated as a cause of BA in Australian livestock, causes EHBD damage in larval zebrafish and mouse EHBD explants [5–7]. Others have reported that mice treated with biliatresone on postnatal day 1 (P1) (80 mg/kg given intraperitoneally) develop a BA-like phenotype [8,9], although prenatal administration to pregnant mothers resulted in either no obvious phenotype (~40-50 mg/kg orally) or prenatal lethality (~40-50 mg/kg intraperitoneally) in their pups [8]. In order to define the relationship between bile acid abnormalities and EHBD damage in BA, we evaluated the histology and liver and serum bile acid profiles of pups born to mothers treated with a low concentration of biliatresone (15mg/kg orally for each of two days).

## Methods

### Animal experiments

BALB/c mice (Jackson Laboratories) were used in this study. All animal experiments were performed in accordance with National Institutes of Health policy and were approved by the Institutional Animal Care and Use Committee at the University of Pennsylvania under protocol # 804862.

Pregnant female mice were administered 15 mg/kg biliatresone or vehicle (containing an equivalent concentration of DMSO, at 0.3 ml/kg) via gavage on days 14 and 15 post mating. Half of the pups from each litter were euthanized at P5 and half at P21. Blood, EHBD, and liver samples were collected from all mothers and pups. Albumin, alkaline phosphatase (ALP), alanine transaminase (ALT) and aspartate transaminase (AST), and bile acids were measured in serum samples. Bile acids and immune cells were measured in liver samples. EHBD and liver samples were also fixed and stained. For details see supplementary data. Since serum sample volumes were limited, not all analyses could be performed on all samples.

### Statistical analysis

Statistical significance was calculated by one and two-tailed Student’s t-tests. Differences in variance were tested using the F test [10]. The number of samples tested for each experiment are given in parentheses in the graphs.

## Results

### Phenotype of pups from biliatresone-treated mothers

P5 and P21 pups from mothers treated with either low-dose biliatresone or vehicle appeared similar with no obvious phenotypic differences. None of the pups developed jaundice or distress or died. There were no histological abnormalities observed for any pups in either liver or bile duct sections (Fig. 1A and C). Stains for HA and collagen I, major components of the extracellular matrix that increase in the setting of fetal/neonatal injury and potential indicators of fibrosis, showed no difference between pups of control and biliatresone-treated mothers, consistent with a lack of damage to the EHBD (Fig. 1B and D) [11]. On average, the two groups of pups had no significant differences in liver biochemistries (Fig. 1C, E), although the treated group at P5 showed significantly higher variability in ALT levels and weight (Fig. 1D).

**Figure 1:**
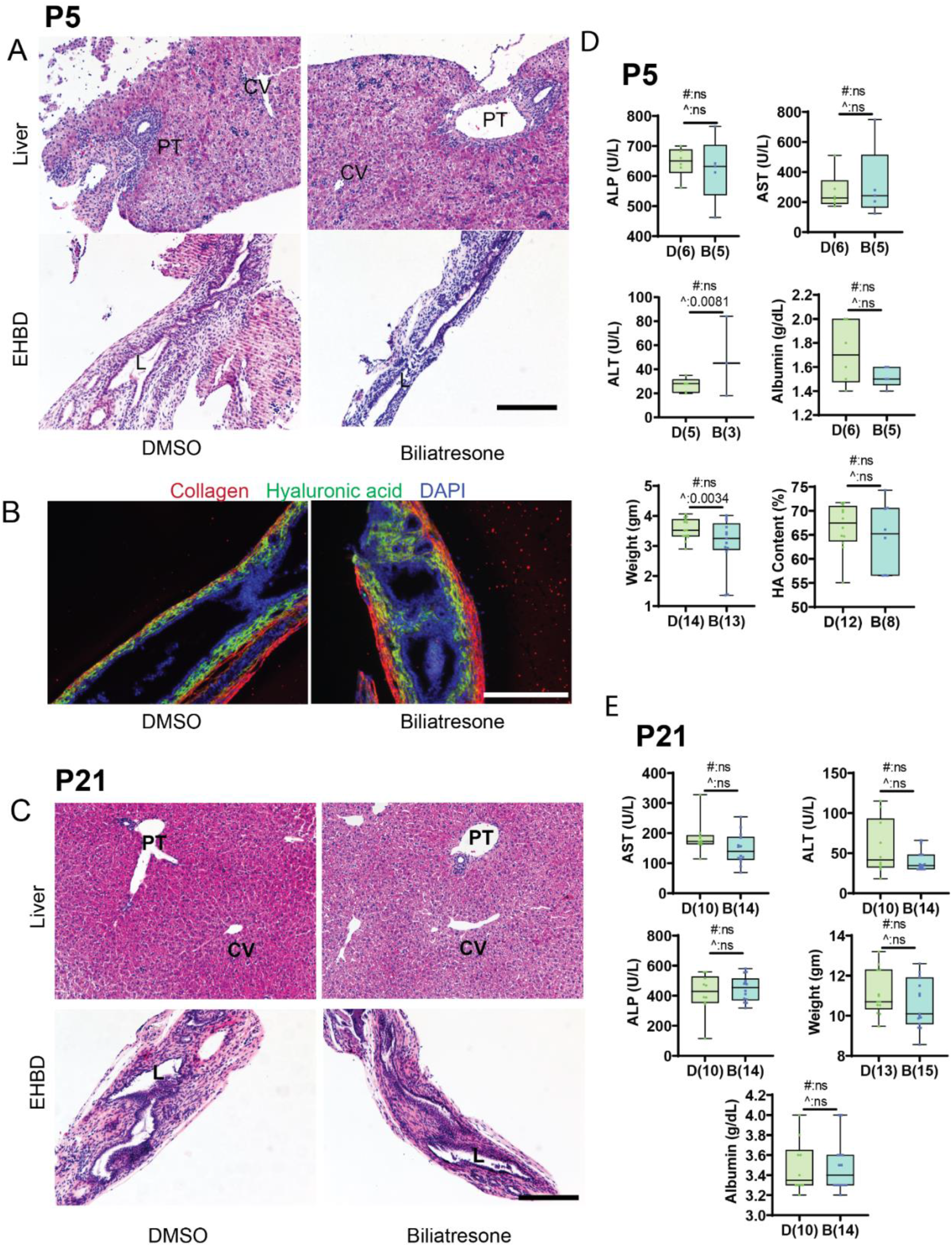
Prenatal biliatresone exposure does not cause significant histological or biochemical differences in pups. (A) H & E staining of liver and EHBD sections isolated from P5 pups born to control and biliatresone-treated mothers. (B) Representative images showing HA (green), collagen (red) and DAPI (blue) staining in P5 pups born to control and biliatresone-treated mothers. (C) H & E staining of liver and EHBD sections isolated from P21 pups born to control and biliatresone-treated mothers. (D) Serum biochemistry and physical parameters of P5 pups. (E) Serum biochemistry and physical parameters of P21 pups. D:DMSO, B:Biliatresone. The number of pups is shown in parentheses. PT: portal triad, CV: central vein, L: lumen. Scale bar, 200 μm. #: t-test p value. ^: F test p value.

### Bile acid profile shows significant differences following biliatresone treatment

There were no significant differences in the liver bile acid profile at P5 (Fig. 2A). However, we did observe a significant increase in serum glycocholic acid (GCA) levels (Fig. 2B). At P21, there was an increase in glycine-conjugated bile acids in both liver and serum. In liver, glycine-β-muricholic acid (GBMCA), glycohyodeoxycholic acid (GHDCA), glycochenodeoxycholic acid (GCDCA) and glycoursodeoxycholic acid (GUDCA) were significantly increased (Fig. 2C), while in serum, GHDCA and GCDCA were increased (Fig. 2D).

**Figure 2:**
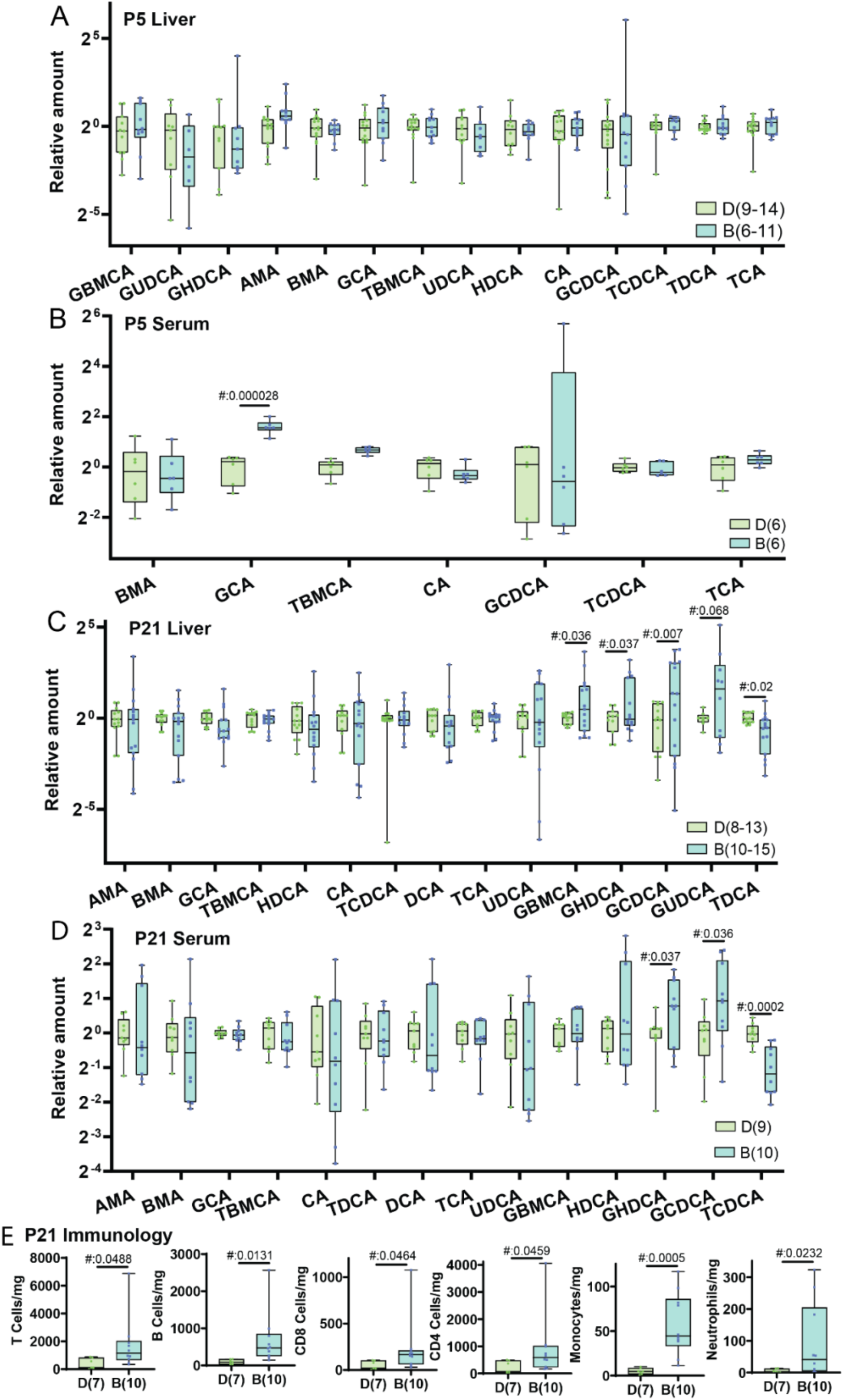
Glycine-modified bile acids and immune infiltrates were increased in pups from biliatresone-treated mothers. (A-D) Relative amounts of various bile acids from liver and serum from P5 and P 21 pups (individual bile acids were normalized to the corresponding mean from control pups). Only bile acids detected in at least 5 samples are shown in the respective graphs. (E) Quantification showing numbers of T-cells, CD4 cells, CD8 cells, B-cells, monocytes and neutrophils in livers isolated from P21 pups. D:DMSO, B: biliatresone, UDCA: ursodeoxycholic acid, HDCA: hyodeoxycholic acid, TDCA: taurodeoxycholic acid, TCA: taurocholic acid, GCDCA: glycochenodeoxycholic acid, CA: cholic acid, TCDCA: taurochenodeoxycholic acid, GCA: glycocholic acid, BMA: β-muricholic acid, TBMCA: tauro-β-muricholic acid, AMA: α-muricholic acid, GHDCA: glycohyodeoxycholic acid, GUDCA: glycoursodeoxycholic acid, DCA: deoxycholic acid and GBMCA: glycine-β-muricholic acid. The number of pups is shown in parentheses. #: t-test p value.

### Significant differences in liver immune profile following biliatresone treatment

Bile acids regulate inflammation, and we thus examined our mouse pups for the presence of inflammatory cells [12]. All P21 animals from biliatresone-treated mothers had significantly elevated liver immune cells compared to control mothers (Fig. 2E). Of the cells identified, B cells (which are required for bile duct injury in the rhesus rotavirus (RRV) model of BA [13]) and monocytes showed the most significant increase.

## Discussion

We demonstrate here that exposing pregnant mice to the biliary toxin biliatresone results in persistent elevations in glycine-conjugated serum bile acids in their pups, without histological evidence of EHBD or liver damage. This suggests that biliatresone can alter bile homeostasis independent of EHBD injury.

Developing a physiologically-relevant BA mouse model based on a prenatal insult has been challenging. This is likely in part because mice are less prone to develop cholestatic diseases than humans [14], but the rarity of BA and the presence of abnormal direct/conjugated bilirubin at birth in many healthy babies who do not develop BA suggests an additional possibility: that the environmental insult responsible for BA causes persistent disease in a minority of cases. If this were the case, finding a BA-afflicted mouse pup even after exposure of the mother to a physiologically-relevant concentration of a toxin or virus might be infrequent [15]. Our aim here was to determine – as a means of understanding the full spectrum of changes in toxic fetal biliary injury – whether exposing mothers to low doses of biliatresone would generate a mild phenotype.

Pups born to control and low-dose biliatresone-treated mothers were phenotypically normal, confirming earlier reports [8], and histological assessment of EHBDs from both groups showed no evidence of damage or recovery post-damage [11]. However, there were several clear differences between the two groups of pups. In the P5 group, two pups had very low weight compared to the average, resulting in a significantly high variance in the biliatresone-treated population. There were also differences in the variance of the ALT levels. This suggests that some animals are more sensitive than others to biliatresone, as was observed for the postnatal biliatresone exposure model developed by Yang et al. and Schmidt et al, which resulted in only a fraction of pups treated with high biliatresone (70-80 mg/kg) developing persistent EHBD damage, with a large fraction of pups showing either no phenotype at all or jaundice followed by recovery [8,9].

The abnormal bile acid metabolites we observed in the pups of biliatresone-treated mouse mothers were similar to those reported for babies with BA. BA patients undergoing operative cholangiography have elevations in multiple liver and plasma bile acids [16].

These abnormalities, observed during the advanced stages of BA, have been attributed to obstruction resulting from bile duct damage. However, Zhou et al [17] found that bile acids (mainly GCA) were elevated in dried blood samples obtained 4 days after birth, before detectable bile duct damage, from infants who later developed BA, suggesting the possibility that abnormal bile acids contribute to or at least precede duct damage. GCA is the major bile acid present in the human neonatal period and it decreases with age; however, in these studies it remained elevated in BA patients [18,19]. Interestingly, our mouse studies identified significant GCA elevations in P5 pups born to mothers administered biliatresone. GCA levels normalized by P21 in mice, but at this age there was a significant increase in other glycine-conjugated bile acids in both liver and serum, especially GHDCA and GCDCA. Glycine modifications are more common in humans than in mice (which favor taurine conjugation), and GCDCA, which results from the modification of CDCA in humans, is the most abundant bile acid in adults [19]; increased GCDCA has been linked to cholestasis and cirrhosis [20]. Since bile acids conjugated to taurine are less toxic than those conjugated to glycine, the low levels of glycine-conjugated bile acids in mice are potentially linked to the low incidence of cholestasis in mice compared to humans. Consistent with this, humanizing mouse bile acids by administering GCDCA leads to increased cholangiocyte injury and collagen deposition, lumen obstruction and sub-epithelial fibrosis in the EHBD [14,21]. Thus, the increase in GCDCA observed in mouse pups after maternal treatment with biliatresone is significant as it suggests the potential for increased bile toxicity, which could cause or exacerbate liver and duct damage. Surprisingly, taurine-conjugated bile acids were not significantly different. Tauro-β-muricholic acid and taurcholic acid were the major taurine-conjugated bile acids, together comprising more than 80 percent of the total bile acid pool.

Bile acids are conjugated to glycine or taurine by bile acid-CoA amino acid N-acyl-transferase (BAAT) enzymes; selectivity is regulated by BAAT selectivity, with different selectivity for a given BAAT found in different species, and by availability of taurine precursors. Mice BAATs are particularly inefficient at conjugating glycine even when taurine availability is reduced [22]. It remains to be seen whether there are any other pathways involved in regulating taurine vs. glycine conjugation. Interestingly, a recent study found that glycine conjugation in mice does not decrease after BAAT knock down, suggesting that alternate pathways are available for glycine conjugation in mice. These could include microbial bile conjugation as well as conjugation mediated by the peroxisomal acyltransferases ACNAT1 and ACNAT2 [23]. It is unknown whether any these pathways are affected by biliatresone exposure.

Because bile acids are known to regulate inflammation [12], we also investigated the immune profile of P21 pups and observed across-the-board increases in immune cells. The results shown here suggest that biliatresone can activate the immune system either independently or by disturbing bile metabolism.

In conclusion, we show that the pups of mice treated with low-dose biliatresone during pregnancy have altered bile acid and immune profiles similar to those observed in BA patients, even in the absence of significant histological changes. Although biliatresone is unlikely to be a cause of human BA, the data suggest that maternal toxin exposure could lead to a spectrum of fetal hepatobiliary injury, with cryptic changes in bile acid modification pathways reflecting mild damage, and that the number of cases of mild injury could markedly outnumber the number progressing to full BA – potentially an explanation for the rarity of the disease.

## Acknowledgements

This study was supported by R01 DK119290 and by a grant from the Fred and Suzanne Biesecker Center for Pediatric Liver Research at the Children’s Hospital of Philadelphia (both to R.G.W.). We are grateful to the NIDDK Center for Molecular Studies in Digestive and Liver Diseases (P30 DK050306) and the UPenn/NIEHS Center of Excellence in Environmental Toxicology for core support (Grant number: P30ES013508).

## Supporting Data

### Materials

The bile acids ursodeoxycholic acid (UDCA), hyodeoxycholic acid (HDCA), taurodeoxycholic acid (TDCA), taurocholic acid (TCA), glycochenodeoxycholic acid (GCDCA), cholic acid (CA), taurochenodeoxycholic acid (TCDCA), glycocholic acid (GCA), β-muricholic acid (BMA), tauro-β-muricholic acid (TBMCA), α-muricholic acid (AMA), glycohyodeoxycholic acid (GHDCA), glycoursodeoxycholic acid (GUDCA), deoxycholic acid (DCA) and glycine-β-muricholic acid (GBMCA) as well as an internal standard stock solution containing a mixture of cholic acid-d4, glycocholic acid-d4 and deoxycholic acid-d4 were obtained from Cayman Chemical (Ann Arbor, Michigan, USA). Stock solutions of 1 mg/mL in 100% methanol were diluted to obtain calibration curves ranging from 15.6 to 1000 nM. Percoll was obtained from GE Healthcare Life Sciences (Chicago, Illinois, USA). Antibodies against CD4, Ly6G and CD90.2 were obtained from BD Pharmingen (San Diego, CA, USA) and antibodies against CD8α, NK1.1, B220 and Ly6C from BioLegend (San Diego, CA, USA). Goat Anti collagen I antibody for staining were obtained from Southern Biotech, Birmingham, AL, USA. Biotinylated hyaluronic acid (HA) binding protein for staining HA were obtained from EMD Millipore, Burlington, MA, USA. LIVE/DEAD fixable viability dye was from Life Technologies (Carlsbad, CA, USA). Biliatresone was synthesized as described [4]. Stock solutions were made in DMSO and diluted in 1X PBS for gavage.

### Histochemistry and immunostaining

EHBDs and livers were formalin-fixed and paraffin embedded, then sectioned to 5 μm. Sections for Hematoxylin and Eosin (H&E) were processed according to standard protocols. Bile duct H&E-stained slides were graded normal or abnormal based on qualitative assessment of lumenal debris, marked inflammation, surface epithelium detachment, and signs of regeneration including multi-layered surface epithelium, peribiliary gland expansion and cholangiocyte hyperplasia. Liver H&E stained slides were graded normal or abnormal based on bile duct expansion and presence of bile plugs and analyzed independently by IDJ and NDT (n>8 animals per group).

For antibody staining, paraffin embedded EHBD sections were deparaffinized with xylene and rehydrated through a graded series of alcohol and distilled water. Antigen retrieval was performed in 10 mM citric acid buffer (pH 6.0). Sections were blocked with 5% bovine serum albumin and permeabilized with 0.4% Triton X-100 prior to antibody incubation. Sections were stained for collagen I, hyaluronic acid (HA) and DAPI as described in [5]. For collagen cy3 anti-goat and for HABP it’s Cy2-straptavidin secondary antibodies were used (1:500, Vector Laboratories).

### Image analysis

Image analysis of stained sections was performed with Fiji ImageJ and QuPath v0.2.0 software. The QuPath selection tool was used to calculate the biliary submucosal area that was occupied by HA relative to the entire submucosal area. As a second measure for the thickness of the HA layer, the width between the lumen and the HA-collagen interface was measured at at-least 5 different places, relative to the entire thickness of the bile duct wall.

### Liver immunology

Intrahepatic leukocytes were isolated by Percoll density gradient centrifugation and stained with LIVE/DEAD fixable viability dye, or with antibodies against CD4, CD8α, NK1.1, B220, Ly6C, Ly6G, and CD90.2. All samples were separated on a MACSQuant flow cytometer (Miltenyi Biotec, Gaithersburg, MD, USA) and analyzed using FlowJo software version 10.6 (Tree Star) (Supporting figure 1).

### Sample processing for HPLC

Bile acids were extracted from homogenized liver and serum samples as described [1,2]. Separations were performed on a Waters BEH C18 Column (2.1 mm x 50 mm 1.7 μm). Mobile phase A was water with 0.1% formic acid, and mobile phase B was methanol with 0.1% formic acid at 0.4 mL/min flow. The gradient started at 5% B and was changed to 40% B over 2 min, then to 99% B over 2 min, held constant for 3 minutes then back to the initial composition for equilibration of the column, for a total chromatographic separation time of 12 min. Analysis was conducted on a Thermo Q Exactive HF coupled to an Ultimate 3000 UHPLC interfaced with a heated electrospray ionization (HESI-II) source. The instrument was operated in negative ion mode alternating between full scan from 250-800 *m/z* at a resolution of 120,000 and parallel reaction monitoring at 60,000 resolution with a precursor isolation window of 0.7 m/z. Since sample amounts were limited, some analysis could not be performed on all samples. Bile acid values were normalized to average values obtained for control pups in each set of experiments. Not all bile acids were detected in the samples; only the bile acids detected were used for analysis.

**Supporting figure 1:**
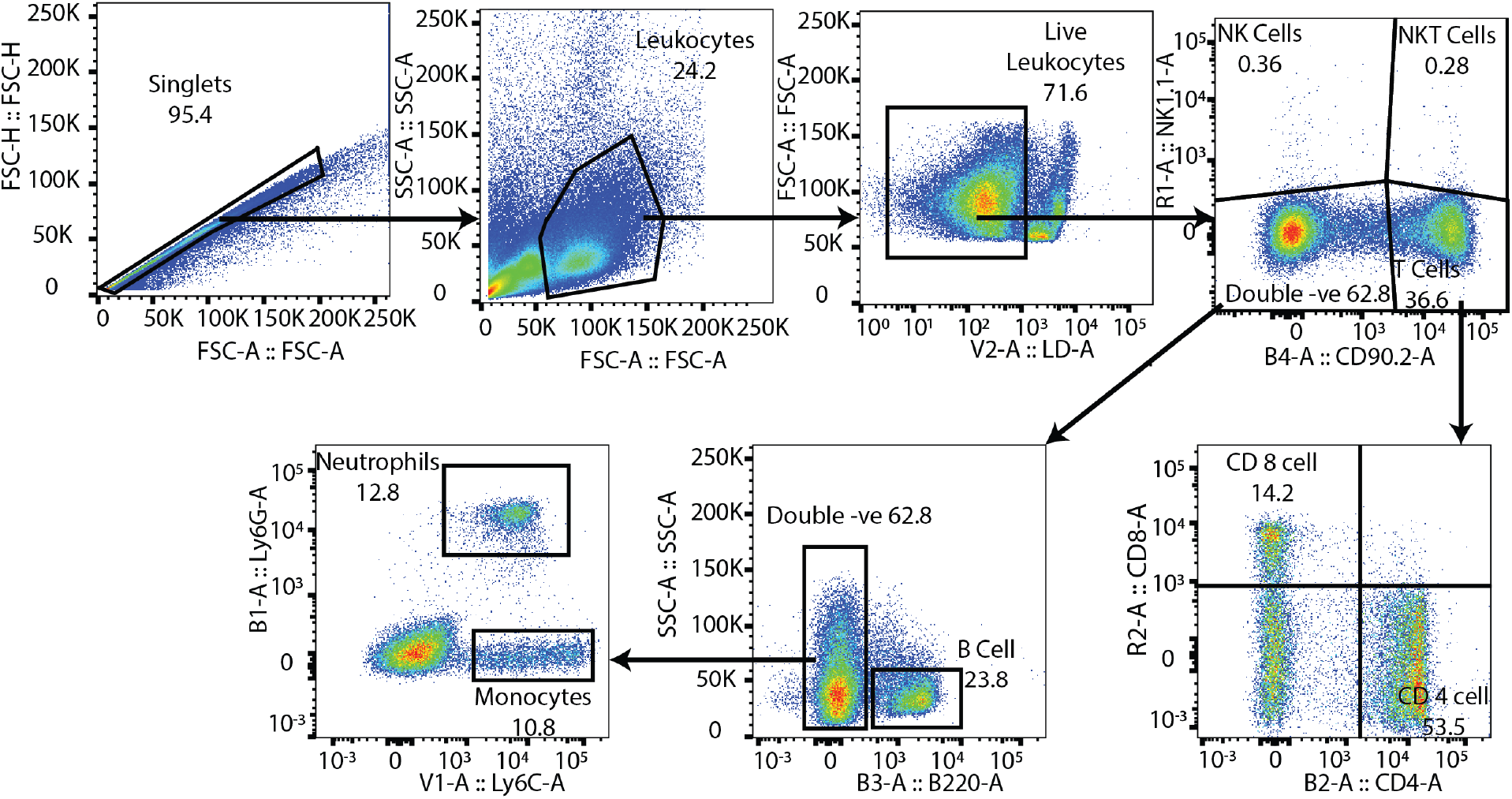
Liver flow gating strategy. Panel used to identify B cells (Live, NK1.1-, CD90.2-,B220+), T-cell populations (Live, NK1.1-, CD90.2+, CD4+ or CD8+), neutrophils (Live, NK1.1-, CD90.2-, B220-, Ly6G+Ly6C+) and monocytes (Live, NK1.1-, CD90.2-, B220-, Ly6G-, Ly6C+).

